# Assessing the performance of real-time epidemic forecasts: A case study of Ebola in the Western Area Region of Sierra Leone, 2014–15

**DOI:** 10.1101/177451

**Authors:** Sebastian Funk, Anton Camacho, Adam J. Kucharski, Rachel Lowe, Rosalind M. Eggo, W. John Edmunds

## Abstract

Real-time forecasts based on mathematical models can inform critical decision-making during infectious disease outbreaks. Yet, epidemic forecasts are rarely evaluated during or after the event, and there is little guidance on the best metrics for assessment. Here, we propose an evaluation approach that disentangles different components of forecasting ability using metrics that separately assess the calibration, sharpness and unbiasedness of forecasts. This makes it possible to assess not just how close a forecast was to reality but also how well uncertainty has been quantified. We used this approach to analyse the performance of weekly forecasts we generated in real time in Western Area, Sierra Leone, during the 2013–16 Ebola epidemic in West Africa. We investigated a range of forecast model variants based on the model fits generated at the time with a semi-mechanistic model, and found that good probabilistic calibration was achievable at short time horizons of one or two weeks ahead but models were increasingly inaccurate at longer forecasting horizons. This suggests that forecasts may have been of good enough quality to inform decision making requiring predictions a few weeks ahead of time but not longer, reflecting the high level of uncertainty in the processes driving the trajectory of the epidemic. Comparing forecasts based on the semi-mechanistic model to simpler null models showed that the best semi-mechanistic model variant performed better than the null models with respect to probabilistic calibration, and that this would have been identified from the earliest stages of the outbreak. As forecasts become a routine part of the toolkit in public health, standards for evaluation of performance will be important for assessing quality and improving credibility of mathematical models, and for elucidating difficulties and trade-offs when aiming to make the most useful and reliable forecasts.

## Introduction

Forecasting the future trajectory of cases during an infectious disease outbreak can make an important contribution to public health and intervention planning. Infectious disease modellers are now routinely asked for predictions in real time during emerging outbreaks (Heesterbeek et al., 2015). Forecasting targets can revolve around expected epidemic duration, size, or peak timing and incidence (Goldstein et al., 2011; Nsoesie et al., 2013; Yang et al., 2015; Dawson et al., 2015), geographical distribution of risk (Lowe et al., 2014), or short-term trends in incidence (Johansson et al., 2016; Liu et al., 2015). However, forecasts made during an outbreak are rarely investigated during or after the event for their accuracy, and only recently have forecasters begun to make results, code, models and data available for retrospective analysis.

The growing importance of infectious disease forecasts is epitomised by the growing number of so-called forecasting challenges. In these, researchers compete in making predictions for a given disease and a given time horizon. Such initiatives are difficult to set up during unexpected outbreaks, and are therefore usually conducted on diseases known to occur seasonally, such as dengue (Johansson et al., 2016; National Oceanic and Atmospheric Administration, 2017; Centres for Disease Control and Prevention, 2017) and influenza (Biggerstaff et al., 2016). The *Ebola Forecasting Challenge* was a notable exception, triggered by the 2013–16 West African Ebola epidemic and set up in June 2015. Since the epidemic had ended in most places at that time, the challenge was based on simulated data designed to mimic the behaviour of the true epidemic instead of real outbreak data. The main lessons learned were that 1) ensemble estimates outperformed all individual models, 2) more accurate data improved the accuracy of forecasts and 3) considering contextual information such as individual-level data and situation reports improved predictions (Viboud et al., 2017).

In theory, infectious disease dynamics should be predictable within the timescale of a single outbreak (Scarpino and Petri, 2017). In practice, however, providing accurate forecasts during emerging epidemics comes with particular challenges such as data quality issues and limited knowledge about the processes driving growth and decline in cases. In particular, uncertainty about human behavioural changes and public health interventions can preclude reliable long-term predictions (Moran et al., 2016; Funk et al., 2017b). Yet, short-term forecasts with an horizon of a few generations of transmission (e.g., a few weeks in the case of Ebola), can yield important information on current and anticipated outbreak behaviour and, consequently, guide immediate decision making.

The most recent example of large-scale outbreak forecasting efforts was during the 2013–16 Ebola epidemic, which vastly exceeded the burden of all previous outbreaks with almost 30,000 reported cases of the disease, resulting in over 10,000 deaths in the three most affected countries: Guinea, Liberia and Sierra Leone. During the epidemic, several research groups provided forecasts or projections at different time points, either by generating scenarios believed plausible, or by fitting models to the available time series and projecting them forward to predict the future trajectory of the outbreak (Fisman et al., 2014; Lewnard et al., 2014; Nishiura and Chowell, 2014; Rivers et al., 2014; Towers et al., 2014; Camacho et al., 2015b; Dong et al., 2015; Drake et al., 2015; Merler et al., 2015; Siettos et al., 2015; White et al., 2015). One forecast that gained attention during the epidemic was published in the summer of 2014, projecting that by early 2015 there might be 1.4 million cases (Meltzer et al., 2014). This number was based on unmitigated growth in the absence of further intervention and proved a gross overestimate, yet it was later highlighted as a “call to arms” that served to trigger the international response that helped avoid the worst-case scenario (Frieden and Damon, 2015). While that was a particularly drastic prediction, most forecasts made during the epidemic were later found to have overestimated the expected number of cases, which provided a case for models that can generate sub-exponential growth trajectories (Chretien et al., 2015; Chowell et al., 2017).

Traditionally, epidemic forecasts are assessed using aggregate metrics such as the mean absolute error (MAE, Chowell, 2017; Pei and Shaman, 2017; Viboud et al., 2017). This, however, only assesses how close the most likely or average predicted outcome is to the true outcome. The ability to correctly forecast uncertainty, and to quantify confidence in a predicted event, is not assessed by such metrics. Appropriate quantification of uncertainty, especially of the likelihood and magnitude of worst case scenarios, is crucial in assessing potential control measures. Methods to assess probabilistic forecasts are now being used in other fields, but are not commonly applied in infectious disease epidemiology (Gneiting and Katzfuss, 2014; Held et al., 2017).

We produced weekly sub-national real-time forecasts during the Ebola epidemic, starting on 28 November 2014. Plots of the forecasts were published on a dedicated web site and updated every time a new set of data were available (Center for the Mathematical Modelling of Infectious Diseases, 2015). They were generated using a model that has, in variations, been used to forecast bed demand during the epidemic in Sierra Leone (Camacho et al., 2015b) and the feasibility of vaccine trials later in the epidemic (Camacho et al., 2015a; Camacho et al., 2017). During the epidemic, we provided sub-national forecasts for the three most affected countries (at the level of counties in Liberia, districts in Sierra Leone and prefectures in Guinea).

Here, we apply assessment metrics that elucidate different properties of forecasts, in particular their probabilistic calibration, sharpness and bias. Using these methods, we retrospectively assess the forecasts we generated for Western Area in Sierra Leone, an area that saw one of the greatest number of cases in the region and where our model informed bed capacity planning.

## Materials and Methods

### Data sources

Numbers of suspected, probable and confirmed Ebola cases at sub-national levels were initially compiled from daily *Situation Reports* (or *SitReps*) provided in PDF format by Ministries of Health of the three affected countries during the epidemic (Camacho et al., 2015b). Data were automatically extracted from tables included in the reports wherever possible and otherwise manually converted by hand to machine-readable format and aggregated into weeks. From 20 November 2014, the World Health Organization (WHO) provided tabulated data on the weekly number of confirmed and probable cases. These were compiled from the patient database, which was continuously cleaned and took into account reclassification of cases avoiding potential double-counting. However, the patient database was updated with substantial delay so that the number of reported cases would typically be underestimated in the weeks leading up to the date of the forecast. Because of this, we used the SitRep data for the most recent weeks until the latest week in which the WHO case counts either equalled or exceeded the SitRep counts. For all earlier times, the WHO data were used.

### Transmission model

We used a semi-mechanistic stochastic model of Ebola transmission described previously (Camacho et al., 2015b; Funk et al., 2017a). Briefly, the model was based on a Susceptible-Exposed-Infectious-Recovered (SEIR) model with fixed incubation period of 9.4 days (WHO Ebola Response Team, 2014), following an Erlang distribution with shape 2. The country-specific infectious period was determined by adding the average delay to hospitalisation to the average time from hospitalisation to death or discharge, weighted by the case-fatality rate. Cases were assumed to be reported with a stochastic time-varying delay. On any given day, this was given by a gamma distribution with mean equal to the country-specific average delay from onset to hospitalisation and standard deviation of 0.1 day. We allowed transmission to vary over time, to capture behavioural changes in the community, public health interventions or other factors affecting transmission for which information was not available at the time. The time-varying transmission rate was modelled using a daily Gaussian random walk with fixed volatility (or standard deviation of the step size) which was estimated as part of the inference procedure (see below). We log-transformed the transmission rate to ensure it remained positive. The behaviour in time can be written as

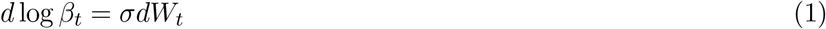

where *β_t_* is the time-varying transmission rate, *W_t_* is the Wiener process and *σ* the volatility of the transmission rate. The basic reproduction number *R*_0,*t*_ at any time was obtained by multiplying *β_t_* with the infectious period. In fitting the model to the time series of cases we extracted posterior predictive samples of trajectories, which we used to generate forecasts.

### Model fitting

Each week, we fitted the model to the available case data leading up to the date of the forecast. Observations were assumed to follow a negative binomial distribution. Since the *ssm* software used to fit the model only implemented a discretised normal observation model, we used a normal approximation of the negative binomial for observations, potentially introducing a bias at small counts. Four parameters were estimated in the process: the initial basic reproduction number *R*_0_ (uniform prior within (1, 5)), initial number of infectious people (uniform prior within (1, 400)), overdispersion of the (negative binomial) observation process (uniform prior within (0, 0.5)) and volatility of the time-varying transmission rate (uniform prior within (0, 0.5)). We confirmed from the posterior distributions of the parameters that these priors did not set any problematic bounds. Samples of the posterior distribution of parameters and state trajectories were extracted using particle Markov chain Monte Carlo (Andrieu et al., 2010) as implemented in the *ssm* library (Dureau et al., 2013). For each forecast, 50,000 samples were extracted and thinned to 5000.

### Predictive model variants

We used the samples of the posterior distribution generated using the Monte Carlo sampler to produce predictive trajectories, using the final values of estimated state trajectories as initial values for the forecasts and simulating the model forward for up to 10 weeks. While all model fits were generated using the same model described above, we tested a range of different predictive model variants to assess the quality of ensuing predictions. We tested variants where trajectories were stochastic (with demographic stochasticity and a noisy reporting process), as well as ones where these sources of noise were removed for predictions. We further tested predictive model variants where the transmission rate continued to follow a random walk (unbounded, on a log-scale), as well as ones where the transmission rate stayed fixed during the forecasting period. Where the transmission rate remained fixed for prediction, we tested variants where we used the final value of the transmission rate and ones where this value would be averaged over a number of weeks leading up to the final fitted point, to reduce the potential influence of the last time point, where the transmission rate may not have been well identified. We tested variants where the predictive trajectory would be based on the final values and start at the last time point, and ones where they would start at the penultimate time point, which could, again, be expected to be better informed by the data. For each model and forecast horizon, we generated point-wise medians and credible intervals from the sample trajectories.

### Null models

To assess the performance of the semi-mechanistic transmission model we compared it to three simpler null models: two representing the constituent parts of the semi-mechanistic model, and a non-mechanistic time series model. For the first null model, we used a *deterministic* model that only contained the mechanistic core of the semi-mechanistic model, that is a deterministic SEIR model with fixed transmission rate and parameters otherwise the same as in the model described elsewhere (Camacho et al., 2015b):

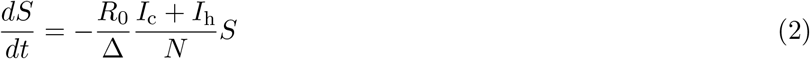

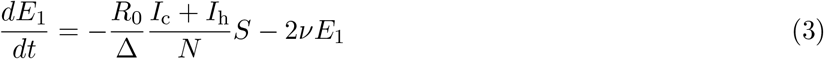

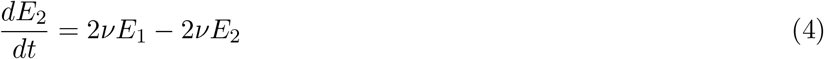

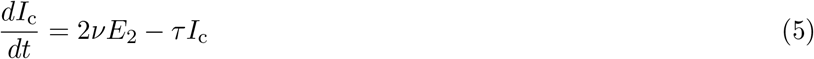

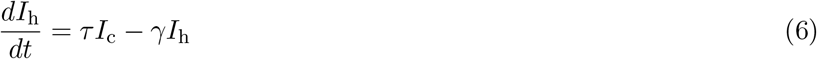

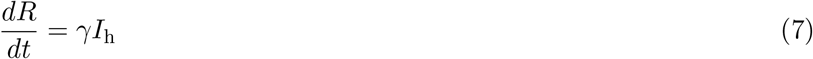

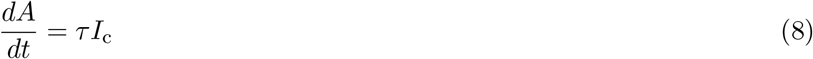

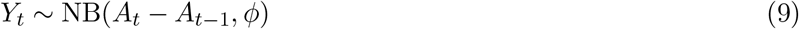

where *Y_t_* are observations at times *t*, *S* is the number susceptible, *E* the number incubating (split into two compartments for Erlang-distributed permanence times with shape 2), *I_c_* is the number infectious and not yet notified, *I*_h_ is the number infectious and notified, *R* is the number recovered or dead, *A* is an accumulator for incidence, *R*_0_ is the basic reproduction number, Δ = 1/*τ* + 1/*ν* is the mean time from onset to outcome, 1/*ν* is the mean incubation period, 1/*τ* + 1/*γ* is the mean duration of infectiousness, 1/*τ* is the mean time from onset to hospitalisation 1/*γ* the mean duration from notification to outcome and NB(*μ*,*φ*) is a negative binomial distribution with mean *μ* and overdispersion *φ*. All these parameters were taken from individual patient observations (WHO Ebola Response Team, 2014) except the overdispersion in reporting *φ*, and the basic reproduction number *R*_0_, which were inferred using Markov-chain Monte Carlo with the same priors as in the semi-mechanistic model.

For the second null model, we used an *unfocused* model where the weekly incidence *Z* itself was modelled using a stochastic volatility model (without drift), that is a daily Gaussian random walk, and forecasts generated assuming the weekly number of new cases was not going to change:

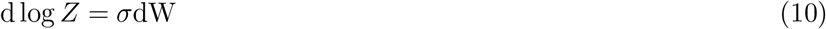

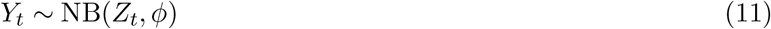

where *Y* are observations, *σ* is the intensity of the random walk and *φ* the overdispersion of reporting (both estimated using Markov-chain Monte Carlo) and dW is the Wiener process.

Lastly, we used a null model based on a non-mechanistic Bayesian autoregressive AR(1) time series model:

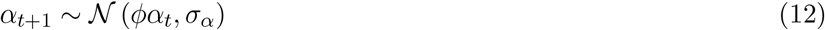

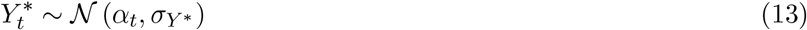

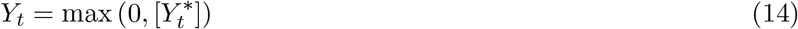

where *φ*, *σ_α_* and *σ*_Y*_ were estimated using Markov-chain Monte Carlo, and […] indicates rounding to the nearest integer. An alternative model with Poisson distributed observations was discarded as it yielded poorer predictive performance.

The deterministic and unfocused models were implemented in *libbi* (Murray, 2015) via the *RBi* (Jacob and Funk, 2017) and *RBi.helpers* (Funk, 2016) *R* packages (R Core Team, 2018). The Bayesian autoregressive time series model was implemented using the *bsts* package (Scott, 2017).

### Metrics

The paradigm for assessing probabilistic forecasts is that they should maximise the sharpness of predictive distributions subject to calibration (Gneiting et al., 2007). We therefore first assessed model calibration at a given forecasting horizon, before assessing their sharpness and other properties.

*Calibration* or reliability (Friederichs and Thorarinsdottir, 2012) of forecasts is the ability of a model to correctly identify its own uncertainty in making predictions. In a model with perfect calibration, the observed data at each time point look as if they came from the predictive probability distribution at that time. Equivalently, one can inspect the probability integral transform of the predictive distribution at time *t* (Dawid, 1984),

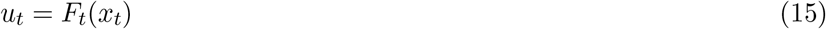

where *x_t_* is the observed data point at time *t* ∈ *t*_1_, …, *t_n_*, *n* being the number of forecasts, and *F_t_* is the (continuous) predictive cumulative probability distribution at time *t*. If the true probability distribution of outcomes at time *t* is *G_t_* then the forecasts *F_t_* are said to be *ideal* if *F_t_* = *G_t_* at all times *t*. In that case, the probabilities *u_t_* are distributed uniformly.

In the case of discrete outcomes such as the incidence counts that were forecast here, the PIT is no longer uniform even when forecasts are ideal. In that case a randomised PIT can be used instead:

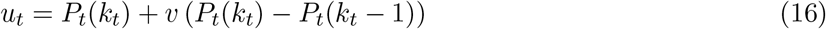

where *k_t_* is the observed count, *P_t_*(*x*) is the predictive cumulative probability of observing incidence *k* at time *t*, *P_t_*(−1) = 0 by definition and v is standard uniform and independent of *k*. If *P_t_* is the true cumulative probability distribution, then *u_t_* is standard uniform (Czado et al., 2009). To assess calibration, we therefore applied the Anderson-Darling test of uniformity to the probabilities *u_t_*. The resulting p-value was a reflection of how compatible the forecasts were with the null hypothesis of uniformity of the PIT, or of the data coming from the predictive probability distribution. We considered that there was no evidence to suggest a forecasting model was miscalibrated if the p-value found was greater than a threshold of *p* ≥ 0.1, some evidence that it was miscalibrated if 0.01 < p < 0.1, and good evidence that it was miscalibrated if p ≤ 0.01. In this context it should be noted, though, that uniformity of the (randomised) PIT is a necessary but not sufficient condition of calibration (Gneiting et al., 2007). The p-values calculated here merely quantify our ability to reject a hypothesis of good calibration, but cannot guarantee that a forecast is calibrated. Because of this, other indicators of forecast quality must be considered when choosing a model for forecasts.

All of the following metrics are evaluated at every single data point. In order to compare the forecast quality of models, they are averaged across the data set.

*Sharpness* is the ability of the model to generate predictions within a narrow range of possible outcomes. It is a data-independent measure, that is, it is purely a feature of the forecasts themselves. To evaluate sharpness at time t, we used the normalised median absolute deviation about the median (MADN) of *y*

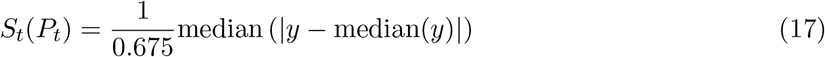

where *y* is a variable with CDF *P_t_*, and division by 0.675 ensures that if the predictive distribution is normal this yields a value equivalent to the standard deviation. The MAD (i.e., the MADN without the normalising factor) is related to the interquartile range (and in the limit of infinite sample size takes twice its value), a common measure of sharpness (Gneiting and Katzfuss, 2014), but is more robust to outliers (Maronna et al., 2018). The sharpest model would focus all forecasts on one point and have *S* = 0, whereas a completely blurred forecast would have *S* → ∞. Again, we used Monte-Carlo samples from *P_t_* to estimate sharpness.

We further assessed the *bias* of forecasts to test whether a model systematically over- or underpredicted. We defined the forecast bias at time *t* as

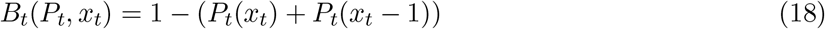

The most unbiased model would have exactly half of predictive probability mass not concentrated on the data itself below the data at time *t* and *B_t_* = 0, whereas a completely biased model would yield either all predictive probability mass above (*B_t_* = 1) or below (*B_t_* = −1) the data.

We further evaluated forecasts using two *strictly proper scoring rules*, that is scores which are minimised if the predictive distribution is the same as the one generating the data. These scores combine the assessment of calibration and sharpness for comparison of overall forecasting skill. The *Ranked Probability Score* (RPS, Epstein, 1969; Murphy, 1969) for count data is defined as (Czado et al., 2009)

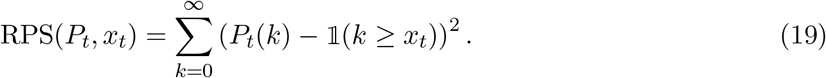

It reduces to the mean absolute error (MAE) if the forecast is deterministic and can therefore be seen as its probabilistic generalisation for discrete forecasts. A convenient equivalent formulation for predictions generated from Monte-Carlo samples is (Gneiting et al., 2007; Czado et al., 2009)

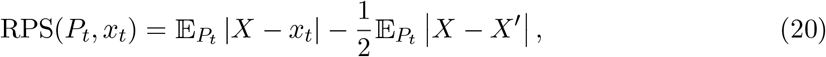

where *X* and *X*′ are independent realisations of a random variable with cumulative distribution *P_t_*.

The *Dawid-Sebastiani score* (DSS) only considers the first two moments of the predictive distribution and is defined as (Czado et al., 2009)

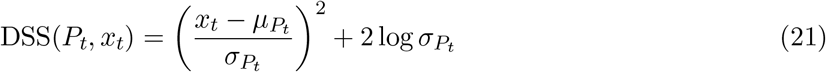

where *μ_P_t__* and *σ*_P_t__ are the mean and standard deviation of the predictive probability distribution, respectively, estimated here using Monte-Carlo samples.

For comparison, we also evaluated forecasts using the *absolute error* (AE) of the median forecast, that is

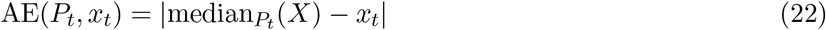

where *X* is a random variable with cumulative distribution *P_t_*.

## Results

The semi-mechanistic model used to generate real-time forecasts during the epidemic was able to reproduce the trajectories up to the date of each forecast, following the data closely by means of the smoothly varying transmission rate (Fig. 1). The overall behaviour of the reproduction number (ignoring depletion of susceptibles which did not play a role at the population level given the relatively small proportion of the population infected) was one of a near-monotonic decline, from a median estimate of 2.9 (interquartile range (IQR) 2.1–4, 90% credible interval (CI) 1.2–6.9) in the first fitted week (beginning 10 August, 2014) to a median estimate of 1.3 (IQR 0.9–1.9, 90% CI 0.4–3.7) in early November, 0.9 (IQR 0.6–1.3, 90% CI 0.2–2.2) in early December, 0.6 in early January (IQR 0.3–0.8, 90% CI 0.1–1.5) and 0.3 at the end of the epidemic in early February (IQR 0.2–0.4, 90% CI 0.1–0.9).

**Figure 1.**
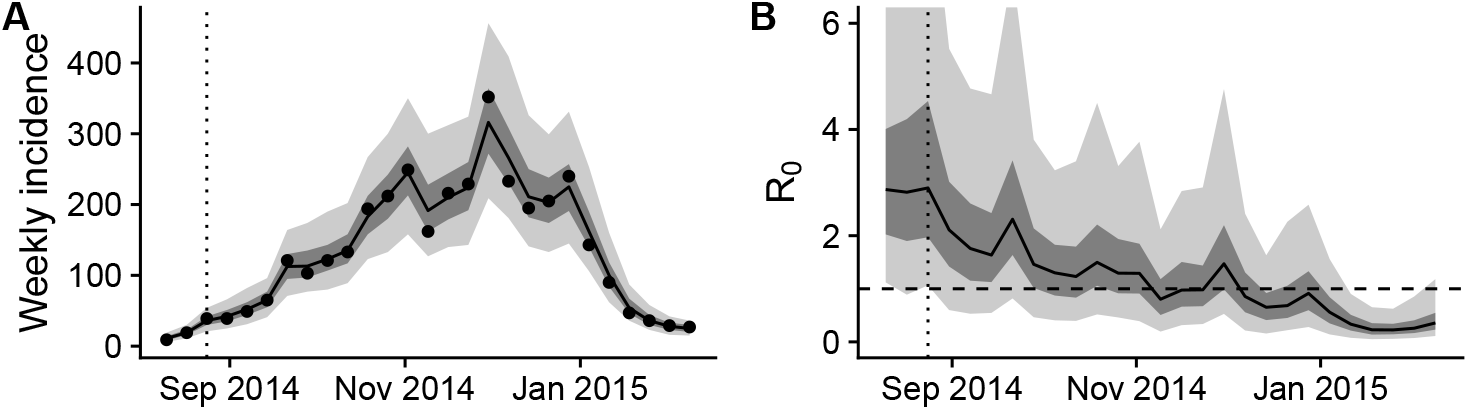
Final fit of the semi-mechanistic model to the Ebola outbreak in Western Area, Sierra Leone. (A) Final fit of the reported weekly incidence (black line and grey shading) to the data (black dots). (B) Corresponding dynamics of the reproduction number (ignoring depletion of susceptibles). Point-wise median state estimates are indicated by a solid line, interquartile ranges by dark shading, and 90% intervals by light shading. The threshold reproduction number (*R*_0_ = 1), determining whether case numbers are expected to increase or decrease, is indicated by a dashed line. In both plots, a dotted vertical line indicates the date of the first forecast assessed in this manuscript (24 August 2014).

The epidemic lasted for a total of 27 weeks, with forecasts generated starting from week 3. For *m*-week ahead forecasts this yielded a sample size of 25 − *m* forecasts to assess calibration. Calibration of forecasts from the semi-mechanistic model were good for a maximum of one or two weeks, but deteriorated rapidly at longer forecasting horizons (Fig. 2). The two semi-mechanistic forecast model variants with best calibration performance used deterministic dynamics starting at the last fitted data point (Table 1). Of these two, the forecast model that kept the transmission rate constant from the value at the last data point performed slightly better across forecast horizons than one that continued to change the transmission rate following a random walk with volatility estimated from the time series. There was no evidence of miscalibration in both of the models with best calibration performance for two-week ahead forecasts, but increasing evidence of mis-calibration for forecast horizons of three weeks or more. Calibration of all model variants was poor four weeks or more ahead, and all the stochastic model variants were miscalibrated for any forecast horizon, including the one we used to publish forecasts during the Ebola epidemic (stochastic, starting at the last data point, no averaging of the transmission rate, no projected volatility).

**Figure 2.**
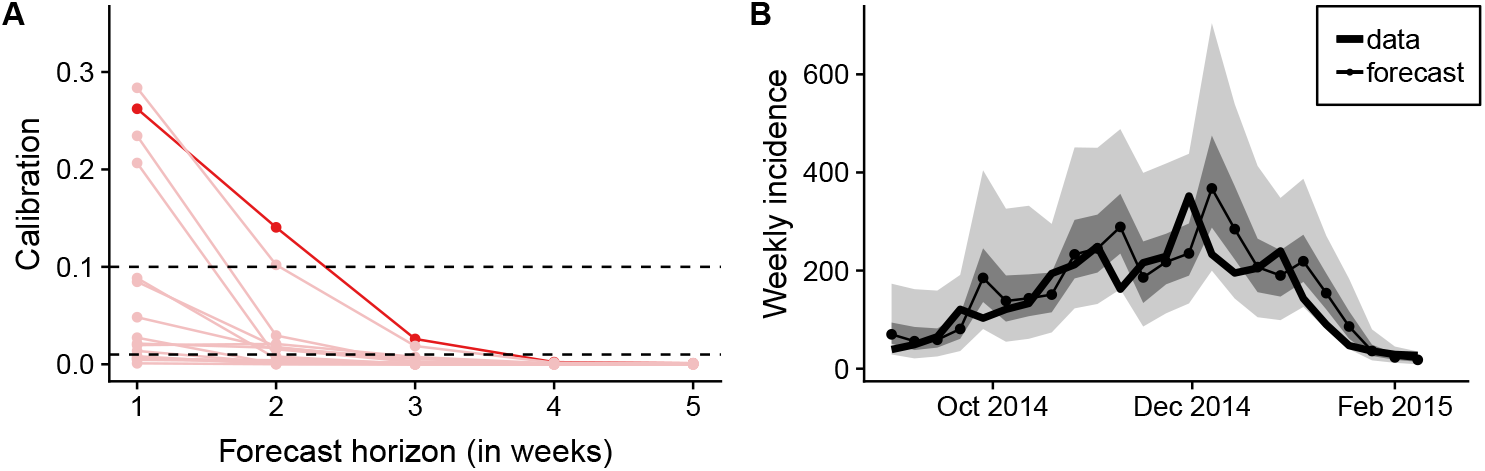
Calibration of forecasts from the semi-mechanistic model. (A) Calibration of model variants (p-value of Anderson-Darling test) as a function of the forecast horizon. Shown in dark red is the best calibrated forecasting model variant (corresponding to the second row of Table 1). Other model variants are shown in light red. (B) Comparison of one-week forecasts of reported weekly incidence generated using the best semi-mechanistic model variant to the subsequently released data. The data are shown as a thick line, and forecasts as dots connected by a thin line. Dark shades of grey indicate the point-wise interquartile range, and lighter shades of grey the point-wise 90% credible interval.

**Table 1.** Calibration of forecast model variants of the semi-mechanistic model. Calibration (p-value of the Anderson-Darling test of uniformity) of deterministic and stochastic forecasts starting either at the last data point or one week before, with varying (according to a Gaussian random walk) or fixed transmission rate either starting from the last value of the transmission rate or from an average over the last 2 or 3 weeks, at different forecast horizons up to 4 weeks. The p-values highlighted in bold reflect predictive models with no evidence of miscalibration. The second row corresponds to the highlighted model in Fig. 2A.

| Predictive model variant |  |  |  | Forecast horizon (weeks) |  |  |  |
| --- | --- | --- | --- | --- | --- | --- | --- |
| stochasticity | start | transmission | averaged | 1 | 2 | 3 | 4 |
| deterministic | at last data point | varying | no | **0.28** | **0.1** | 0.02 | <0.01 |
| deterministic | at last data point | fixed | no | **0.26** | **0.14** | 0.03 | <0.01 |
| deterministic | at last data point | fixed | 2 weeks | **0.24** | 0.03 | <0.01 | <0.01 |
| deterministic | at last data point | fixed | 3 weeks | **0.21** | <0.01 | <0.01 | <0.01 |
| deterministic | 1 week before | varying | no | 0.05 | 0.02 | <0.01 | <0.01 |
| deterministic | 1 week before | fixed | no | 0.09 | 0.02 | <0.01 | <0.01 |
| deterministic | 1 week before | fixed | 2 weeks | 0.09 | <0.01 | <0.01 | <0.01 |
| deterministic | 1 week before | fixed | 3 weeks | 0.03 | <0.01 | <0.01 | <0.01 |
| stochastic | at last data point | varying | no | 0.02 | 0.02 | <0.01 | <0.01 |
| stochastic | at last data point | fixed | no | 0.02 | 0.02 | <0.01 | <0.01 |
| stochastic | at last data point | fixed | 2 weeks | 0.01 | <0.01 | <0.01 | <0.01 |
| stochastic | at last data point | fixed | 3 weeks | <0.01 | <0.01 | <0.01 | <0.01 |
| stochastic | 1 week before | varying | no | <0.01 | <0.01 | <0.01 | <0.01 |
| stochastic | 1 week before | fixed | no | <0.01 | <0.01 | <0.01 | <0.01 |
| stochastic | 1 week before | fixed | 2 weeks | <0.01 | <0.01 | <0.01 | <0.01 |
| stochastic | 1 week before | fixed | 3 weeks | <0.01 | <0.01 | <0.01 | <0.01 |

The calibration of the best semi-mechanistic forecast model variant (deterministic dynamics, transmission rate fixed and starting at the last data point) was better than for any of the null models (Fig. 3A and Table 2) for up to three weeks. While there was no evidence for miscalibration of the autoregressive null model for 1-week-ahead forecasts, there was good evidence of miscalibration for longer forecast horizons. There was some evidence of miscalibration of the unfocused null model, which assumes that the same number of cases will be reported in the weeks following the week during which the forecast was made, for 1 week ahead and good evidence of miscalibration beyond. Calibration of the deterministic null model was poor for all forecast horizons.

**Figure 3.**
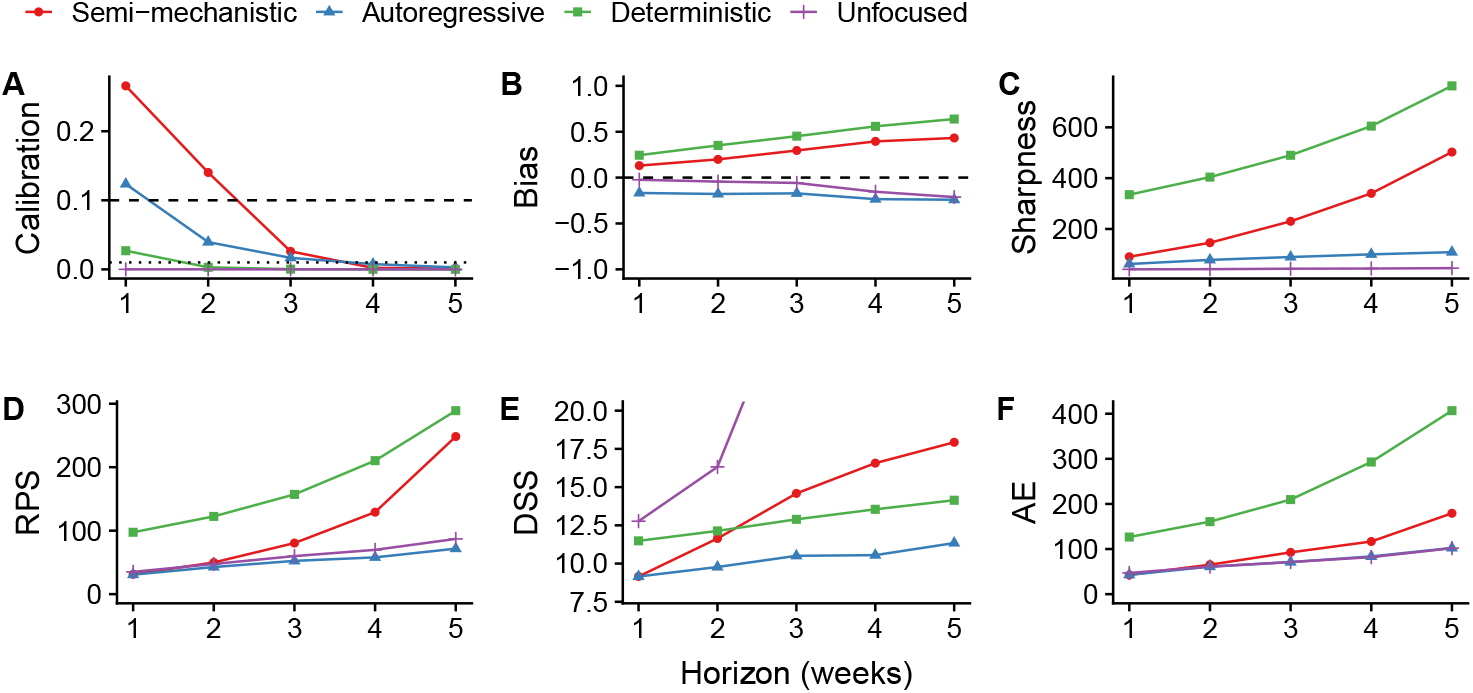
Forecasting metrics and scores of the best semi-mechanistic model variant compared to null models. Metrics shown are (A) calibration (p-value of Anderson-Darling test, greater values indicating better calibration, dashed lines at 0.1 and 0.01), (B) bias (less bias if closer to 0), (C) sharpness (MAD, sharper models having values closer to 0), (D) RPS (better values closer to 0), (E) DSS (better values closer to 0) and (F) AE (better values closer to 0), all as a function of the forecast horizon.

**Table 2.** Forecasting metrics and scores of the best semi-mechanistic model variant compared to null models. The values shown are the same scores as in Fig. 3, for forecasting horizons up to three weeks. The p-values for calibration highlighted in bold reflect predictive models with no evidence of miscalibration.

| Model | Calibration | Sharpness | Bias | RPS | DSS | AE |
| --- | --- | --- | --- | --- | --- | --- |
| **1 week ahead** |  |  |  |  |  |  |
| Semi-mechanistic | **0.26** | 91 | 0.13 | 31 | 9.2 | 42 |
| Autoregressive | **0.1** | 61 | -0.17 | 31 | 9.1 | 43 |
| Deterministic | 0.03 | 340 | 0.24 | 97 | 11 | 130 |
| Unfocused | <0.01 | 41 | -0.024 | 35 | 13 | 47 |
| **2 weeks ahead** |  |  |  |  |  |  |
| Semi-mechanistic | **0.14** | 150 | 0.2 | 50 | 12 | 65 |
| Autoregressive | 0.03 | 77 | -0.18 | 43 | 9.9 | 60 |
| Deterministic | <0.01 | 400 | 0.35 | 120 | 12 | 160 |
| Unfocused | <0.01 | 42 | -0.044 | 48 | 16 | 61 |
| **3 weeks ahead** |  |  |  |  |  |  |
| Semi-mechanistic | 0.03 | 230 | 0.3 | 81 | 15 | 93 |
| Autoregressive | 0.02 | 90 | -0.17 | 53 | 11 | 73 |
| Deterministic | <0.01 | 490 | 0.45 | 160 | 13 | 210 |
| Unfocused | <0.01 | 44 | -0.058 | 60 | 29 | 71 |

The semi-mechanistic and deterministic models showed a tendency to overestimate the predicted number of cases, while the autoregressive and null models tended to underestimate (Fig. 3B and and Table 2). This bias increased with longer forecast horizons in all cases. The semi-mechanistic model with best calibration progressed from a 12% bias at 1 week ahead to 20% (2 weeks), 30% (3 weeks), 40% (4 weeks) and 44% (5 weeks) overestimation. At the same time, this model showed rapidly decreasing sharpness as the forecast horizon increased (Fig. 3C and and Table 2). This is reflected in the proper scoring rules that combine calibration and sharpness, with smaller values indicating better forecasts (Fig. 3D-E and and Table 2). At 1-week ahead, the mean RPS values of the autoregressive, unfocused and best semi-mechanistic forecasting models were all around 30. At increasing forecasting horizon, the RPS of the semi-mechanistic model grew faster than the RPS of the autoregressive and unfocused null models. The DSS of the semi-mechanistic model, on the other hand, was very similar to the one of the autoregressive and better than that of the other null models at a forecast horizon of 1 week, with the autoregressive again performing best at increasing forecast horizons.

Focusing purely on the median forecast (and thus ignoring both calibration and sharpness), the absolute error (AE, Fig.3F and Table 2) was lowest (42) for the best semi-mechanistic model variant at 1-week ahead forecasts, although similar to the autoregressive and unfocused null models. With increasing forecasting horizon, the absolute error increased at a faster rate than for the autoregressive and unfocused null models.

We lastly studied the calibration behaviour of the models over time; that is, using the data and forecasts available up to different time points during the epidemic (Fig. 4). This shows that from very early on, not much changed in the ranking of the different semi-mechanistic model variants. Comparing the best semi-mechanistic forecasting model to the null models, again, for almost the whole duration of the epidemic calibration of the semi-mechanistic model was best for forecasts 1 or 2 weeks ahead.

**Figure 4.**
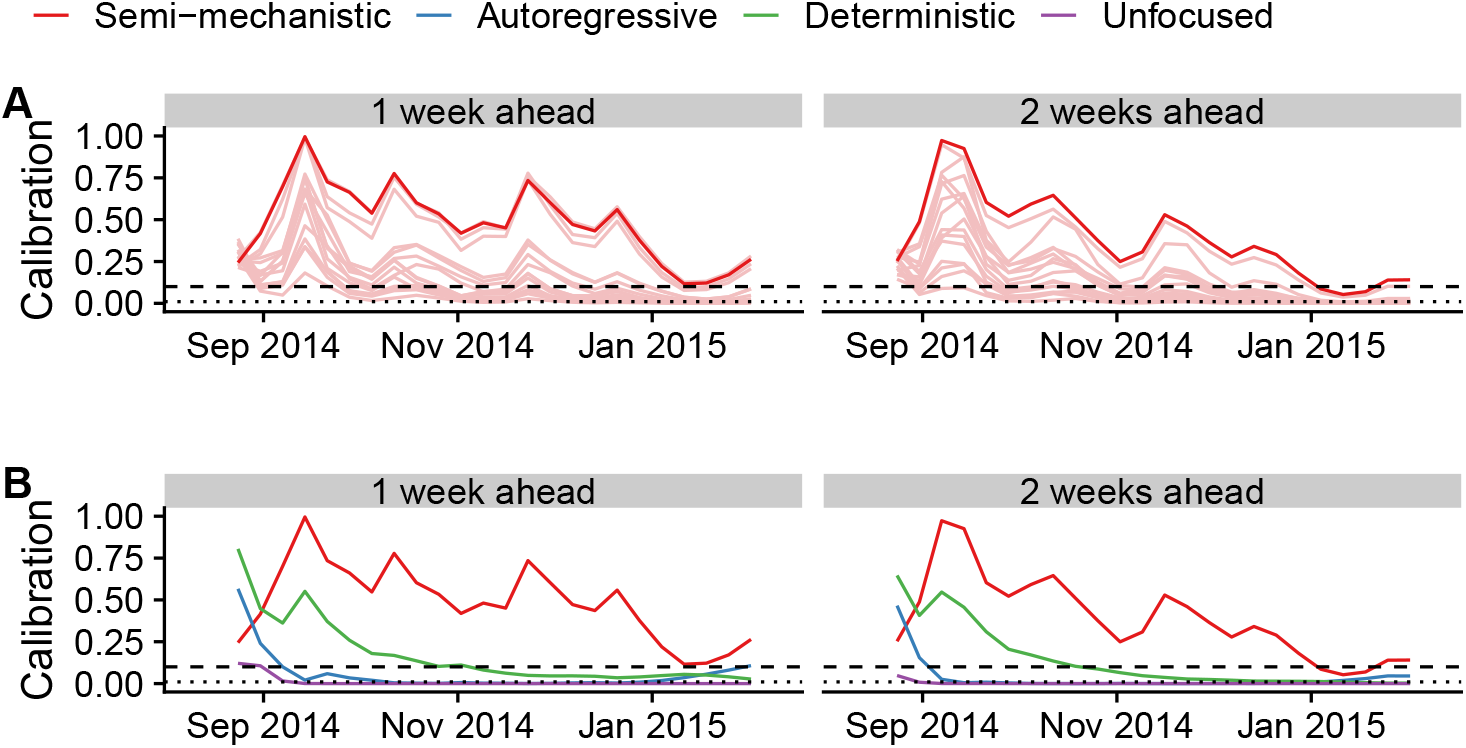
Calibration over time. Calibration scores of the forecast up to the time point shown on the x-axis. (A) Semi-mechanistic model variants, with the best model highlighted in dark red and other model variants are shown in light red. (B) Best semi-mechanistic model and null models. In both cases, 1-week (left) and 2-week (right) calibration (p-value of Anderson-Darling test) are shown.

## Discussion

Probabilistic forecasts aim to quantify the inherent uncertainty in predicting the future. In the context of infectious disease outbreaks, they allow the forecaster to go beyond merely providing the most likely future scenario and quantify how likely that scenario is to occur compared to other possible scenarios. While correctly quantifying uncertainty in predicted trajectories has not commonly been the focus in infectious disease forecasting, it can have enormous practical implications for public health planning. Especially during acute outbreaks, decisions are often made based on so-called “worst-case scenarios” and their likelihood of occurring. The ability to adequately assess the magnitude as well as the probability of such scenarios requires accuracy at the tails of the predictive distribution, in other words good calibration of the forecasts.

Probabilistic forecasts need to be assessed using metrics that go beyond the simple difference between the central forecast and what really happened. Applying a suite of assessment methods to the forecasts we produced for Western Area, Sierra Leone, we found that probabilistic calibration of semi-mechanistic model variants varied, with the best ones showing good calibration for up to 2–3 weeks ahead, but performance deteriorated rapidly as the forecasting horizon increased. This reflects our lack of knowledge about the underlying processes shaping the epidemic at the time, from public health interventions by numerous national and international agencies to changes in individual and community behaviour. During the epidemic, we only published forecasts up to 3 weeks ahead, as longer forecasting horizons were not considered appropriate.

Our forecasts suffered from bias that worsened as the forecasting horizon expanded. Generally, the forecasts tended to overestimate the number of cases to be expected in the following weeks, as did most other forecasts generated during the outbreak (Chretien et al., 2015). This is in line with previous findings where our model was applied to predict simulated data of a hypothetical Ebola outbreak (Funk et al., 2017a). Log-transforming the transmission rate in order to ensure positivity skewed the underlying distribution and made very high values possible. Moreover, we did not model a trend in the transmission rate, whereas in reality transmission decreased over the course of the epidemic, probably due to a combination of factors ranging from better provision of isolation beds to increasing awareness of the outbreak and subsequent behavioural changes. While our model captured changes in the transmission rate in model fits, it did not forecast any trends such as the observed decrease over time. Capturing such trends and modelling the underlying causes would be an important future improvement of real-time infectious disease models used for forecasting.

There are trade-offs between achieving good outcomes for the different forecast metrics we used. Deciding whether the best forecast is the best calibrated, the sharpest or the least biased, or some compromise between the three, is not a straightforward task. Our assessment of forecasts using separate metrics for probabilistic calibration, sharpness and bias highlights the underlying trade-offs. While the best calibrated semi-mechanistic model variant showed better calibration performance than the null models, this came at the expense of a decrease in the sharpness of forecasts. Comparing the models using the RPS alone, the semi-mechanistic model of best calibration performance would not necessarily have been chosen. Following the paradigm of maximising sharpness subject to calibration, we therefore recommend to treat probabilistic calibration as a prerequisite to the use of forecasts, in line with what has recently been suggested for post-processing of forecasts (Wilks, 2018). Probabilistic calibration is essential for making meaningful probabilistic statements (such as the chances of seeing the number of cases exceed a set threshold in the upcoming weeks) that enable realistic assessments of resource demand, the possible future course of the epidemic including worst-case scenarios, as well as the potential impact of public health measures. Once calibration is ensured, other criteria such as the RPS or DSS can be used to select the best model for forecasts, or to generate weights for ensemble forecasts combining several models. Such ensemble forecasts have become a standard in weather forecasting (Gneiting and Raftery, 2005) and have more recently shown promise for infectious disease forecasts (Yamana et al., 2016; Yamana et al., 2017; Viboud et al., 2017).

Other models may have performed better than the ones presented here. Because we did not have access to data that would have allowed us to assess the importance of different transmission routes (burials, hospitals and the community) we relied on a relatively simple, flexible model. The deterministic SEIR model we used as a null model performed poorly on all forecasting scores, and failed to capture the downturn of the epidemic in Western Area. On the other hand, a well-calibrated mechanistic model that accounts for all relevant dynamic factors and external influences could, in principle, have been used to predict the behaviour of the epidemic reliably and precisely. Yet, lack of detailed data on transmission routes and risk factors precluded the parameterisation of such a model and are likely to do so again in future epidemics in resource-poor settings. Future work in this area will need to determine the main sources of forecasting error, whether structural, observational or parametric, as well as strategies to reduce such errors (Pei and Shaman, 2017).

In practice, there might be considerations beyond performance when choosing a model for forecasting. Our model combined a mechanistic core (the SEIR model) with non-mechanistic variable elements. By using a flexible non-parametric form of the time-varying transmission rate, the model provided a good fit to the case series despite a high levels of uncertainty about the underlying process. Having a model with a mechanistic core came with the advantage of enabling the assessment of interventions just as with a traditional mechanistic model. For example, the impact of a vaccine could be modelled by moving individuals from the susceptible into the recovered compartment (Camacho et al., 2015a; Camacho et al., 2017). At the same time, the model was flexible enough to visually fit a wide variety of time series, and this flexibility might mask underlying misspecifications. More generally, when choosing between forecast performance and the ability to explicitly account for the impact of interventions, a model that accounts for the latter might, in some cases, be preferable. Where possible, the guiding principle in assessing real-time models and predictions for public health should be the quality of the recommended decisions based on the model results (Probert et al., 2018).

Epidemic forecasts played a prominent role in the response to and public awareness of the Ebola epidemic (Frieden and Damon, 2015). Forecasts have been used for vaccine trial planning against Zika virus (World Health Organization, 2017) and will be called upon again to inform the response to the next emerging epidemic or pandemic threat. Recent advances in computational and statistical methods now make it possible to fit models in near-real time, as demonstrated by our weekly forecasts (Center for the Mathematical Modelling of Infectious Diseases, 2015). Such repeated forecasts are a prerequisite for the use of metrics that assess not only how close the predictions were to reality, but also how well uncertainty in the predictions has been quantified. An agreement on standards of forecast assessment is urgently needed in infectious disease epidemiology, and retrospective or even realtime assessment of forecasts should become standard for epidemic forecasts to prove accuracy and improve end-user trust. The metrics we have used here or variations thereof could become measures of forecasting performance that are routinely used to evaluate and improve forecasts during epidemics.

For forecast assessment to happen in practice, evaluation strategies must be planned before the forecasts are generated. In order for such evaluation to be performed retrospectively, all forecasts as well as the data, code and models they were based on should be made public at the time, or at least preserved and decisions recorded for later analysis. We published weekly updated aggregate graphs and numbers during the Ebola epidemic, yet for full transparency it would have been preferable to allow individuals to download raw forecasts for further analysis.

If forecasts are not only produced but also evaluated in real time, this can give valuable insights into strengths, limitations, and reasonable time horizons. In our case, by tracking the performance of our forecasts, we would have noticed the poor calibration of the model variant chosen for the forecasts presented to the public, and instead selected better calibrated variants. At the same time, we did not store the predictive distribution samples for any area apart from Western Area in order to better use available storage space, and because we did not deem such storage valuable at the time. This has precluded a broader investigation of the performance of our forecasts.

At the same time, research into modelling and forecasting methodology and predictive performance at times during which there is no public health emergency should be part of pandemic preparedness activities. To facilitate this, outbreak data must be made available openly and rapidly. Where available, combination of multiple sources, such as epidemiological and genetic data, could increase predictive power. It is only on the basis of systematic and careful assessment of forecast performance during and after the event that predictive ability of computational models can be improved and lessons be learned to maximise their utility in future epidemics.

